# Micromechanics of lung capillaries across mouse lifespan and in positive- vs negative- pressure ventilation

**DOI:** 10.1101/2025.03.02.641015

**Authors:** Kathryn Regan, Lauren Castle, Robert LeBourdais, Abdulrahman Kobayter, Linzheng Shi, Winita Wangsrikhun, Gabrielle Grifno, Rohin Banerji, Athanasios Batgidis, Belá Suki, Hadi T. Nia

## Abstract

The lung undergoes continuous remodeling throughout normal development and aging, including changes to alveolar and capillary structure and function. While histological methods allow static analysis of these age-related changes, characterizing the changes that occur in response to mechanical stimuli remains difficult, particularly over a dynamic, physiologically relevant range in a functioning lung. Alveolar and capillary distension – the change in diameter of alveoli and capillaries, respectively, in response to pressure changes – is one such process, where dynamically controlling and monitoring the diameter of the same capillary or alveolus is essential to infer its mechanical properties. We overcome these limitations by utilizing the recently developed crystal ribcage to image the alveoli and vasculature of a functional mouse lung across the lifespan in postnatal (6-7 days), young adult (12-18 weeks), and aged (20+ months) mice. Using a range of biologically relevant vascular (0-15 cmH_2_O) and transpulmonary (3-12 cmH_2_O) pressures, we directly quantify vascular and alveolar distention in the functional lung as we precisely adjust pulmonary pressures. Our results show differences in age-related alveolar and vascular distensibility: when we increase transpulmonary alveolar or vascular pressure, vessels in postnatal lungs expand less and undergo less radial and axial strain, under each respective pressure type, suggesting stiffer capillaries than in older lungs. However, while vessels in young adult and aged lungs respond similarly to variations in vascular pressure, differences in elasticity start to emerge at the alveolar scale in response to transpulmonary alveolar pressure changes. Our results further indicate that differing effects of ventilation mode (i.e., positive vs negative) present themselves at the capillary level, with vessels under positive pressure undergoing more compression than when under negative-pressure conditions. These findings contribute both to the understanding of the functional changes that occur within the lung across the lifespan, as well as to the debate of ventilation effects on lung microphysiology.

## Introduction

Throughout the course of the normal lifespan, the structural and mechanical properties of the lung undergo progressive remodeling^1–6^, consequently altering the physics, biology, and immunology of the lung^7^. Given the delicate interplay between structure and function in the lung^8,9^, these changes impact key physiological processes such as gas exchange, immune defense, and disease progression^10,11^ which can, in turn, influence additional remodeling^12^. At the microscale, understanding how alveoli and capillaries respond to mechanical inputs such as pressure changes is vital for understanding the overall functionality of the lung in health^13–15^, disease^16,17^, and aging.

Alveolar and capillary distension, or the way that alveoli and capillaries change diameter, is one such pressure-related measure that provides additional insight into the stiffness of the alveoli and capillaries. Recent work found that circulating cancer cells undergo pressure-dependent stresses within the lung vasculature during the breathing cycle, the magnitude of which depended on the local vessel diameter: cells in smaller capillaries experienced a higher degree of stress as pressure was increased^18^. Additionally, given the highly mechanosensitive behavior of immune cell populations of the lung^19,20^, the role of mechanosensing in innate immunity^21^, and the hematopoietic nature of the lung^22^, alveolar and capillary distensibility could be a valuable focus in understanding cancer and immune cell trafficking, sequestration, and activation. Indeed, any age-dependent changes to distensibility could have implications on immune response and overall health.

A growing body of literature has documented age-related changes in tissue structure and mechanics using static techniques such as atomic force microscopy (AFM) and histology^2,4^, yet these endpoint methods fail to capture the dynamic nature and stiffness anisotropy of intact, functional tissue. Both histology and AFM require invasive tissue preparation, compromising the unique air-liquid interface of the lung tissue. Additionally, AFM only measures the stiffness perpendicular to the cut surface, thus failing to capture the stiffness anisotropy in the radial and axial directions of the capillaries. Small animal models enable characterization of the whole tissue responses to pressure and ventilation, as well as changes in elasticity, tissue stiffness, and lung compliance in situ across the lifespan^2,23^, but high-resolution, dynamic imaging at the capillary and alveolar scales remains limited. Outside of experimental quantification, mathematical and computational tools can model blood flow and circulatory dynamics in the lung^24^, and have been used to couple age-related mechanical changes to experimentally-observed tissue strain, breathing work, and alveolar dynamics^25,26^. However, the limited, often static nature of experimental boundary conditions constrains the depth of information they can supply. Characterizing the pressure-induced transpulmonary and vascular responses that occur in a functioning lung across the lifespan thus remains an ongoing question.

In addition to development, illness, and aging, the mode of ventilation (i.e., positive-pressure vs negative-pressure ventilation) is believed to have a significant impact on the lung’s mechanical and structural properties. While mechanical ventilation can treat severe respiratory illnesses, it can also result in ventilator induced lung injury (VILI) ^27–29^. Thus, several studies have probed the benefits and risks of positive-pressure ventilation^25,30–32^, as in mechanical ventilation, in comparison to negative-pressure ventilation^33,34^, as in spontaneous breathing. The resulting literature is divided, with some studies claiming no physiological difference exists between the two ventilation schemes^23^ while several others emphasize the beneficial features of negative-pressure ventilation, citing increased oxygenation and decreases in VILI^33,35^. While these studies primarily focus on respiratory function of the lung, they lack insight into how ventilation mode within the same lung affects circulation at the capillary level.

To address this knowledge gap present at the alveolar and capillary level, we utilize the recently developed crystal ribcage^8^, an optically transparent shell that recapitulates the murine ribcage geometry and enables high-resolution imaging of the intact, functional murine lung at alveolar and capillary level resolution. Using the crystal ribcage, we investigate how alveolar and capillary distensibility changes across the murine lifespan. Combining our experimental findings with a mathematical model, we also estimate axial and radial Young’s moduli of the capillaries. We further investigate the effect of positive- versus negative-pressure ventilation on the murine vasculature at both the alveolar and capillary levels. This work represents a valuable measure of the age-related micromechanical changes in respiration-circulation coupling that occur across the lifespan, with direct impact on the ongoing scientific debate on the advantages and disadvantages of positive- vs negative-pressure ventilation.

## Results

### An experimental method for measuring alveolar and capillary distension across the murine lifespan using the crystal ribcage

The crystal ribcage (Fig. 1A) enables independent and precise control over both the pressure difference between the alveoli and pleural surface (the transpulmonary pressure; applied by positive- or negative-pressure ventilation) and the pressure at the capillaries (the vascular pressure). The crystal ribcage additionally possesses the ability to non-invasively switch between positive- and negative-pressure ventilation within the same lung. These pressure controls allow for precise control and high-resolution imaging of the lung (Fig. 1B). To study alveolar and capillary distension across different age groups, we designed three ribcages tailored for developing postnatal (6-7 days), young adult (12-18 weeks), and aged (20+ months) C57BL/6 mice (Fig. 1C). We utilized transgenic reporter mice whose cells express tdTomato as a structural label for the postnatal and young adult groups, and we perfused in Evans Blue as a vascular label for all the age groups. Imaging both fluorescent channels, we obtained high-resolution images of alveoli, capillaries (tdTomato), and vessel lumens (Evans Blue) (Fig 1D(i)), enabling direct assessment of how the tissue distends as pressures are varied (Fig. 1D(ii)). This innovative platform allows for the direct investigation of alveolar level micromechanics and vascular distension throughout the murine lifespan under controllable, biologically relevant conditions.

**Figure 1.**
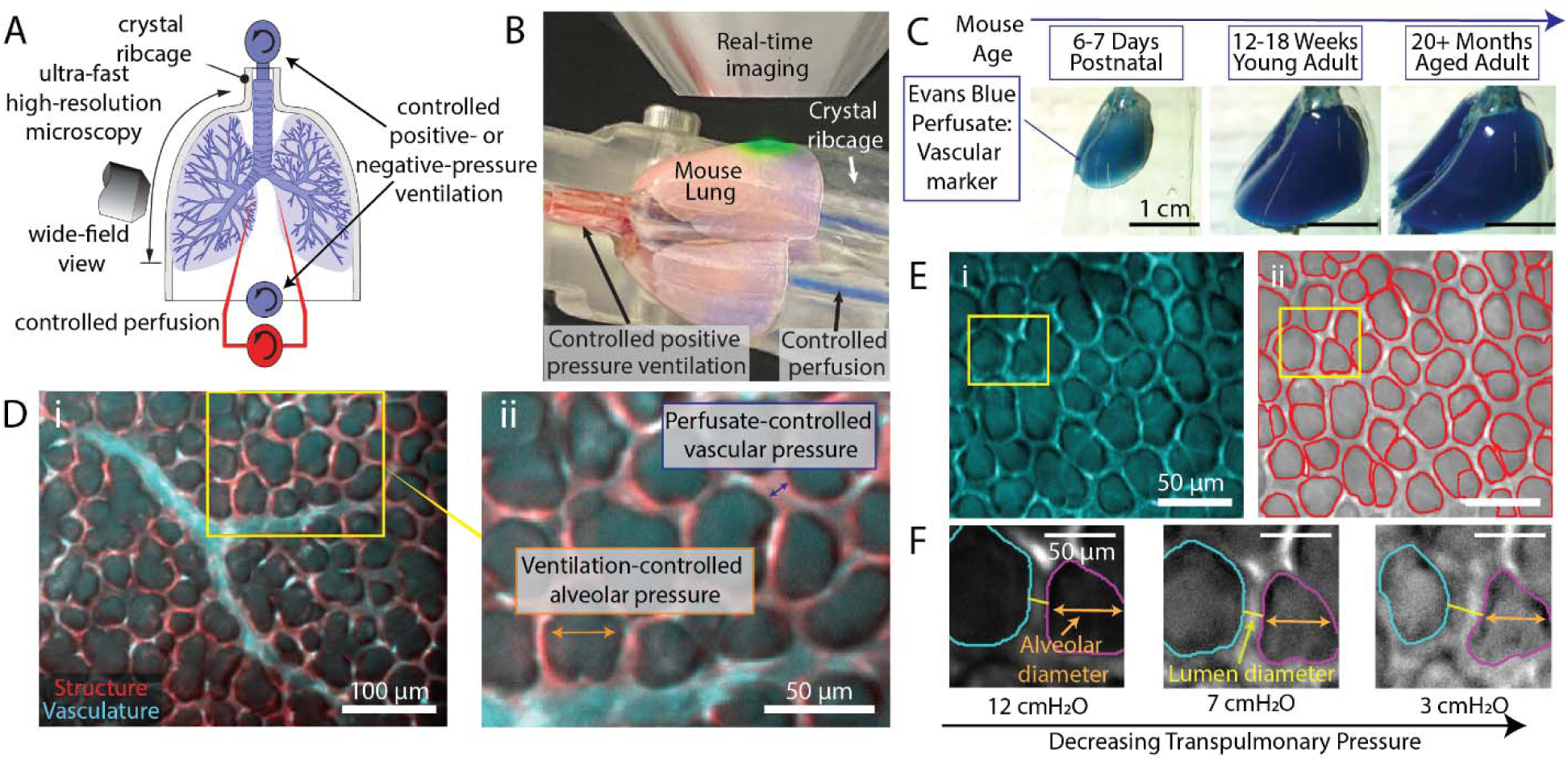
Varying vascular and alveolar pressures to quantify alveolar and vascular distention across the murine lifespan in the crystal ribcage. (A) Schematic representation of the crystal ribcage, a transparent ribcage that conforms to the surface of the murine lung to recreate the native ribcage geometry, and that retains the ability to control ventilation (either positive- or negative-pressure) and perfusion during high-resolution microscopy. (B) Confocal imaging enables real-time imaging of the tissue while controlling both vascular and transpulmonary pressure. (C) We measure alveolar and capillary distensibility across the lifespan using age-specific crystal ribcages for postnatal (6-7 days), young adult (12-18 weeks), and aged (20+ months) mice. Inset images show sample lungs post-perfusion with Evans Blue vascular label. (D) Transgenic reporter mice possess an intrinsic fluorescent structure label (red) while perfusion of Evans Blue dye (cyan) labels the vascular lumen (i) allowing for the quantification of alveolar and lumen diameters under a variety of physiological pressure changes (ii, inset from (i)) within the same lung at the same region of interest. (E) Confocal images (i) taken in the same lung at the same ROI at each pressure are segmented (ii) using a generative artificial intelligence-powered automated segmentation algorithm. The image shown in (i) was taken under positive pressure conditions with a transpulmonary pressure of 7 cmH_2_O and vascular pressure of 0 cmH_2_O. (F) Custom-written Python scripts match alveoli and lumens between the different pressures, enabling the measurement of individual diameters across the pressure range. All images shown in (F) were taken under positive-pressure conditions.

A key advantage of the crystal ribcage is that it allows users to image the same region of interest across the pressure ranges, thereby minimizing errors associated with mouse-to-mouse variability and enabling precise characterization of the dynamics of the individual structures (i.e., individual alveoli, capillaries, and vascular lumens). We combine high-resolution images of paired alveoli and lumen with a custom-trained AI-based analysis pipeline^36^ to determine how the diameter of individual alveoli and lumen respond to increasing pressure. (Fig. 1E, 1F, Supplemental Figure 1). We then fed the lumen diameters into a mathematical model for calculating the radial and axial Young’s modulus of capillaries across a range of pressures, derived such as to account for the strain stiffening that occurs within the lung. This approach is thus able to capture the mechanical properties of tissue at the alveolar and capillary level.

### Nonlinear age dependent trends in capillary distention in response to increasing vascular pressure

We control vascular pressure at the whole lung scale by increasing the pressure at the pulmonary artery while clamping closed the tube connected to the pulmonary vein. The pressure across the capillaries then rises and generates an outward, radial force on the vessel lumens (Fig. 2A). To ensure the perfusate pressure at the capillaries is equal to the pressure measured at the pulmonary artery, we chose to change the pressure quasi-statically, i.e., wait until the pressure equilibrates after each change of pressure, before each imaging step. While the quasistatic change in pressure may not fully capture the dynamics of pressure change in circulation-respiration, it remains the only approach that allows for a direct estimation of pressure in capillaries and alveoli. To isolate only the role of vascular pressure in capillary distension, we hold the transpulmonary pressure constant at 7 cmH_2_O while applying biologically relevant quasistatic vascular pressures ranging from 0-15 cmH_2_O (Fig. 2A); within the vasculature, 15 cmH_2_O approximates the mean vascular pressure experienced at the mouse pulmonary artery^37^. We conducted high-resolution imaging at each pressure to quantify the mechanical response of the lumen to applied forces (Fig. 2B, Supplemental Figure 2). We previously reported that increasing vascular pressure causes an increase in vascular diameter in young mice^8^ but, to our knowledge, capillary micromechanics across the murine lifespan have not been reported. Representative confocal images show clear differences in structure between postnatal (Fig. 2B, left), young adult (Fig 2B, middle), and aged (Fig. 2B, right) lumens. Even at low vascular pressures (Fig 2B(i)), vessel lumens in postnatal mice were notably larger than young adult and aged mice. Upon increasing the vascular pressure from 0 to 15 cmH_2_O (Fig. 2B(ii)), vessels distended as the increased internal pressure of the vessel exerted radial forces on the lumen walls.

**Figure 2.**
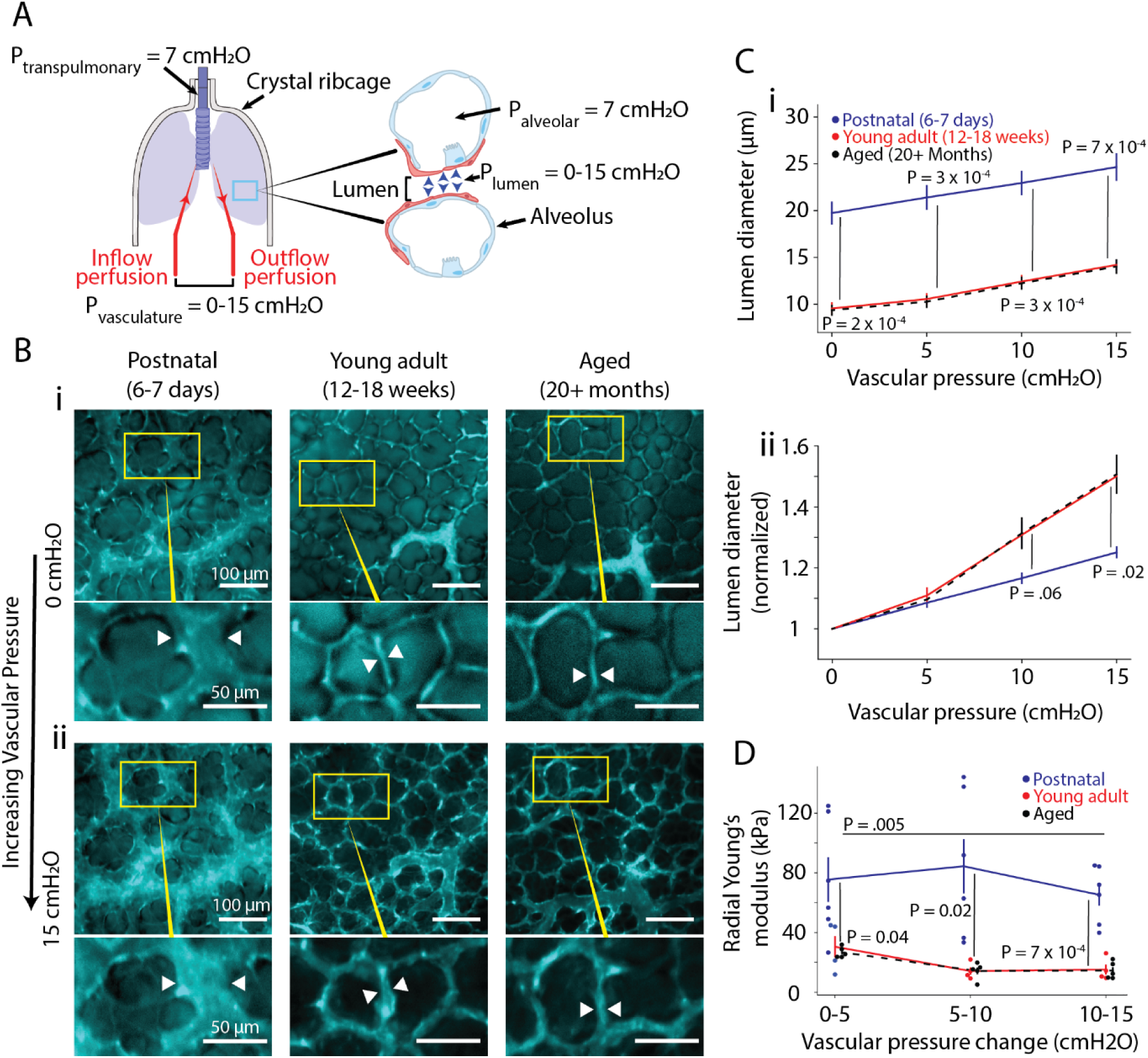
Stiffer aged capillaries display reduced expansion in response to increasing vascular pressure. (A). While transpulmonary pressure is maintained at 7 cmH_2_O via tracheal cannulation, we apply varying vascular pressures ranging from 0 to 15 cmH_2_O. Sensors at inflow and outflow points verify that pressure is equal across the capillary bed. (B) Confocal images capture changes in the diameters of vessel lumens in response to vascular pressure changes. Evans Blue dye labels the vessel lumens via vascular perfusate. Shown are representative images of postnatal (left), young adult (middle), and aged (right) vessels at vascular pressures of 0 (i) and 15 (ii) cmH_2_O. Insets (yellow) show how individual vessels respond to the change in pressure. As the pressure increases, the diameter of the lumen (white arrowheads) increases. Postnatal vessels (left) undergo smaller changes in response to the pressure change than vessels in young adult (middle) and aged (right) mice. (C) Lumens are statistically larger in postnatal mice at all vascular pressures measured (i) and experienced a smaller relative change in diameter per unit pressure increase (ii). Mathematical modeling uses experimentally measured lumen diameters to calculate a radial Young’s modulus for vessels; postnatal lungs have significantly higher radial stiffness than aged or young adult lungs at all pressures (D). While young adult lungs remain statistically similar across all pressures (blue curve), aged lungs show a significantly stiffer radial Young’s modulus at low (0-5 cmH_2_O) pressures than at higher (5- 10cmH_2_O, 10-15cmH_2_O) pressures (p = 0.005). Plots show mean + standard error for N = 4 mice for each condition. Statistics are determined by two-tailed t-test; due to the high degree of overlap in young and aged data, when statistical comparisons between groups are shown they represent postnatal vs young adult and young adult vs aged.

Lumen diameter measurements across the pressure range reflect these structural differences (Fig. 2C(i)). The mean diameter at 0 cmH_2_O was found to be 19.74 + 3.26 µm in postnatal lungs, compared to 9.55 + 1.53 µm in young adult and 9.37 + 1.48 µm in aged lungs. Lumen diameter in all age groups increased as vascular pressure increased, yet young adult and aged samples did not approach postnatal sizes and in fact stayed highly aligned. These findings suggest substantially lower vascular resistance to flow in postnatal lungs, potentially reflecting developmental adaptations to improve blood flow and oxygenation in the developing lung.

To quantify age-related differences in capillary distensibility, we normalized the diameter measurements to the diameter at 0 cmH_2_O (Fig. 2C(ii)). By doing so, we focus on the rate of change relative to the unstressed state and reveal notable differences in the compliance of the lumens. Intriguingly, lumens at all ages initially exhibited similar behavior at low pressures, as evidenced in the overlap of normalized data between 0 and 5 cmH_2_O: the normalized lumen diameter at 5 cmH_2_O lie within a range of 0.03 normalized units across all ages. However, in the range of 5 cmH_2_O to 15 cmH_2_O the stiffness of postnatal lungs diverges from young adult and aged lungs, with normalized lumen diameters at 15 cmH_2_0 of 1.25 + 0.05 versus 1.5 + 0.12 and 1.51 + 0.16 in postnatal, young adult, and aged mice respectively.

Next, we coupled the experimental measurements of lumen diameter with mathematical modeling to estimate the Young’s modulus of the vessels in the radial (outward) direction (Fig. 2D). To capture the effect of native strain stiffening phenomena, we characterize three Young’s modulus values for the incremental differences between the four vascular pressure points. As expected, we find that postnatal lungs have significantly larger radial Young’s modulus values at all pressure ranges (> 65 kPa), remaining statistically similar across the pressure ranges measured. Young adult and aged lungs exhibit significantly lower radial Young’s modulus values; interestingly, while both ages follow similar trends in decreasing radial Young’s modulus with increasing vascular pressure ranges, only aged lungs are significantly stiffer at the lowest (0-5 cmH_2_O) pressure range than at higher ranges. As aging is typically assumed to stiffen vessels, these unexpectedly aligned radial elasticity values represent a previously under-characterized trait, highlighting distinct lumen behaviors and shedding light on intriguing structural and functional changes across the lifespan.

### Postnatal lumens are axially stiffer than those of young adult and aged mice

Next, we investigated how alveoli and lumen across the murine lifespan responded to increasing positive transpulmonary pressure. We imaged the lung at biologically relevant transpulmonary pressures between 3 and 12 cmH_2_O, varying the transpulmonary pressure by applying a static positive pressure at the trachea and preserving the vascular pressure at 0 cmH_2_O. Under positive-pressure ventilation conditions, a positive pressure is applied at the trachea that generates a positive pressure gradient and forces air into the lungs. This pressure causes an increase in intra-alveolar pressure, a radial stretch of the alveolar walls, and a corresponding axial stretch of the vascular lumen (Fig. 3A). This expansion of alveoli is critical for oxygen exchange, as their ability to expand and contract facilitates the transport of oxygen and carbon dioxide into and out of the lung.

**Figure 3.**
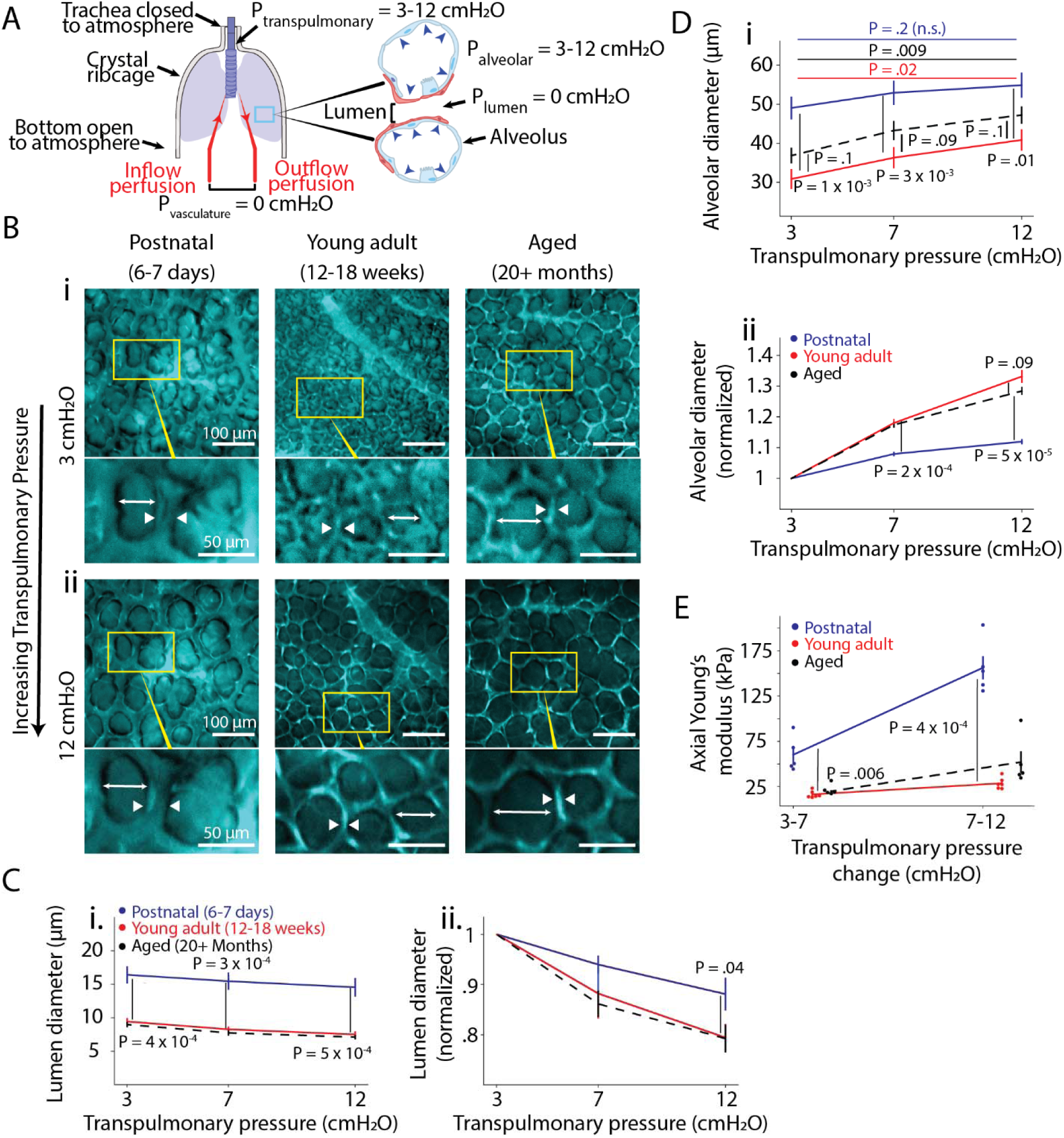
Age dependent differential alveolar and lumen expansion due to transpulmonary pressure changes. (A) Under positive-pressure ventilation, a positive gradient of air pressure is applied across the lung via the trachea. The transpulmonary pressure at the whole lung scale is varied between 3-12 cmH_2_O, and the pressure within the vasculature is held constant at 0 cmH_2_O. At the alveolar scale, the pressure within the alveolus is equal to the transpulmonary pressure and longitudinally stretches the alveolar wall to equilibrate along the pressure gradient as the lung expands. (B) Confocal microscopy images of the vessels perfused with Evans Blue dye depict the alveolar scale response to increasing air pressure in postnatal (left), young adult (middle), and aged (right) lungs. At lower (i) transpulmonary pressures, diameters of alveoli (white arrow) are smaller than at higher (ii) transpulmonary pressures. In contrast lumens (white arrowheads) are larger at lower pressures than higher. (C) Postnatal lungs have larger lumens across the pressure range (i) and display higher stiffnesses when normalized to their lowest pressure value in comparison to young adult and aged lumens (ii). (D) Postnatal lungs have larger alveoli across the observed pressure ranges (i) and have a smaller relative change in alveolar diameter across the pressure range in comparison to young adult and aged lungs (ii), implying stiffer alveolar walls than in their older counterparts. (E) Mathematical modeling to estimate the axial Young’s modulus of the capillaries found that postnatal lungs are significantly stiffer in the axial direction, with significant increases in axial Young’s Modulus at higher pressures. Plots present mean + standard error over N = 5 (postnatal), N = 6 (young adult), and N = 5 (aged) mice. Statistics are determined by two-tailed t-test; due to the high degree of overlap in young and aged data, when statistical comparisons between groups are shown they represent postnatal vs young adult and young adult vs aged.

Using Evans Blue dye to label the lumen of the lung vasculature, we compared alveolar and lumen diameter changes across age groups as we varied the transpulmonary pressure (Fig. 3B, Supplemental Fig. 3) with imaging regions paired at low (Fig. 3B(i)) and high (Fig. 3B(ii)) pressures. Postnatal lungs possessed significantly larger lumens compared to young adult and aged at all pressures measured (Fig. 3C(i)). Consistent with previous findings^8^, we found that lumen diameters decreased as transpulmonary pressure increased, stretching the interpenetrating vessel lumens under an axial strain. Upon normalization to 3 cmH_2_O, the pressure-induced decreases in lumen diameter were consistent in young adult and aged lungs, and larger than in postnatal lumens (Fig. 3C(ii)). The smaller change in normalized lumen diameter of postnatal lungs suggests that they possess higher radial stiffnesses in response to the stretching forces exerted by positive alveolar pressure.

Alveolar diameters showed an inverse relationship with lumen diameters, increasing in diameter as lumens decreased in a manner consistent with previous findings^8^. Postnatal lungs possessed larger alveoli compared to young adult and aged mice (Fig. 3D(i)). Alveoli in aged mice trended towards having larger diameters than in young adults, while both ages exhibited significant increases in diameter over the experimental pressure range (Fig. 3D(i)). These results agree with prior studies^2,5^, confirming that murine lungs undergo significant structural changes throughout the lifespan.

We then normalized the alveolar diameters to the average values at 3 cmH_2_O, quantifying the rate of change in alveolar diameter, a measure which describes the stiffness of the alveolar septum as it stretches along the axial direction during alveolar inflation. Postnatal alveoli had significantly smaller changes in normalized alveolar diameter, suggesting a higher axial stiffness compared to young adult and aged counterparts (Fig. 3D(ii)). Aged alveoli possessed a smaller normalized diameter than younger alveoli, trending towards significance for normalized values (Fig 3. D(ii)). These results suggest an intriguing nonlinear relationship with alveolar wall stiffness, with postnatal lungs being more resistant to changes in transpulmonary pressure, with this resistance first decreasing, then increasing over the lifespan.

We confirm these stiffness approximations with mathematical modeling to calculate the axial Young’s modulus of the septal wall (Fig. 3E). Our modeling captures the alveolar strain stiffening that occurs with pressure changes and finds that the axial stiffness of postnatal lungs increases almost threefold across the experimental pressure range. These values are more than twice the axial Young’s modulus of young adult and aged alveolar walls: while both ages have similar Young’s modulus values between 3 and 7 cmH_2_O, aged lungs are significantly stiffer at the higher (7-12 cmH_2_O) pressure change. These values are comparable to other recently reported values for alveolar wall stiffness of 12 and 15 kPa in young adult and aged mice, respectively, likely corresponding to our low pressure regime^38^. Overall, despite the structural changes that occur in both lumen and alveoli throughout aging, only alveolar responses exhibit an evolving relationship to increasing transpulmonary pressure across the murine lifespan as highlighted by their differences in axial Young’s modulus.

### Positive-pressure ventilation results in higher capillary distension compared to negative- pressure ventilation

Next, we sought to compare how alveolar and capillary distension varies between positive- and negative-pressure conditions, as in mechanical ventilation and spontaneous breathing, respectively. In our setup, under negative pressure the trachea is open to atmosphere (0 cmH_2_O) and the lung inflates and deflates following negative pressure gradients induced in the closed chest cavity. Thus, the transpulmonary pressure is greater than zero at the whole lung scale yet the pressure within the alveolus remains at or close to 0 cmH_2_O, with slight fluctuations around 0 cmH_2_O arising from air flow in the lung (Fig. 4A). Unlike in positive- pressure ventilation, there is no outward deforming force on the alveoli or vessels: instead, the alveoli and vessels expand and contract due to the stretching and relaxing of the lung that occurs as negative pressure is applied to the chest cavity.

**Figure 4.**
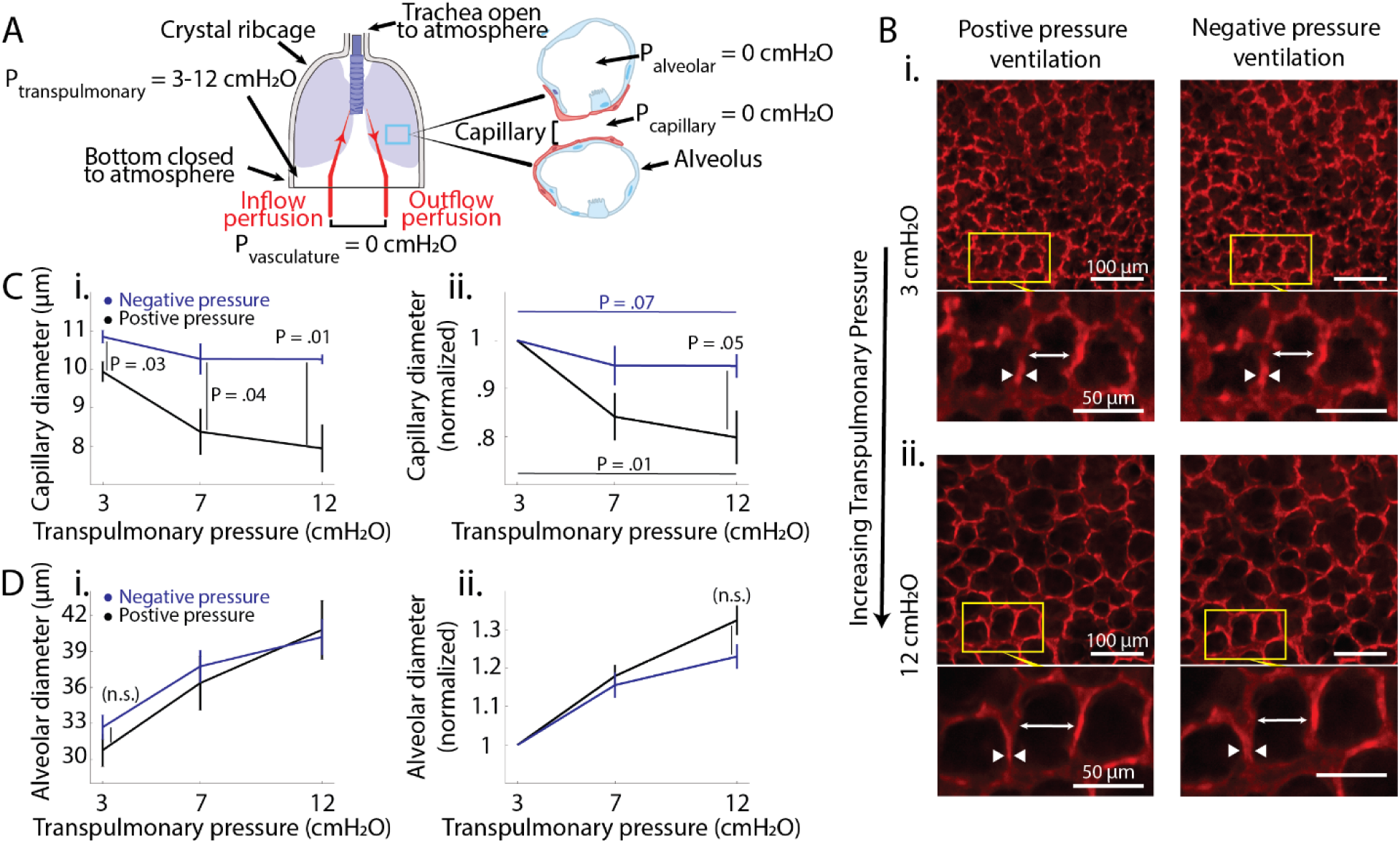
Capillaries of young adult mice exhibit reduced diameter change under negative pressure compared to positive pressure. (A) Under negative-pressure ventilation, the trachea is open to air while negative pressure is applied to the sealed base of the crystal ribcage. At the whole lung scale, transpulmonary pressure is varied from 3-12 cmH_2_O and can be increased and decreased to ventilate the lung; however, air pressure at the alveolus is constant at approximately 0 cmH_2_O. Vascular pressure is maintained at 0 cmH_2_O. (B) Confocal microscopy images show the alveolar scale response to increasing air pressure under both positive-pressure ventilation (left), whereby a positive air pressure gradient is exerted via the trachea, and negative pressure ventilation (right) as described in (A). All images are taken at the same ROI within the same lung at transpulmonary pressure values ranging from 3 cmH_2_O (i) to 12 cmH_2_O (ii). As the pressure increases, the diameters of the alveoli (white arrow) increase while the diameters of the capillaries (white arrowheads) decrease. (C) Capillaries are larger (i) and show a smaller relative change in diameter (ii) under negative-pressure conditions compared to positive-pressure conditions. (D) The absolute alveolar diameter (i) and normalized diameter change (ii) due to increased transpulmonary pressure is similar in both positive- and negative-pressure ventilation. Plots present mean + standard error over N = 4 mice. Statistics are determined by two-tailed t-test.

Using young adult (12-18 week) transgenic reporter mice with structure labeled in tdTomato, we imaged the same ROIs under both positive and negative pressure at the same values of transpulmonary pressures (3-12 cmH_2_O), using confocal microscopy to capture the capillary and alveolar structures at low (Fig. 4B(i)) and elevated (Fig. 4B(ii)) pressures. Significant differences in capillary diameter emerged between these transpulmonary pressure modes. While capillaries under both conditions compressed in the radial direction with increasing alveolar pressure, capillaries were significantly larger under negative pressure than under positive pressure (Fig. 4C(i)), with the difference increasing as the pressure rose. These differences in capillary diameter indicate that the resistance to flow is notably higher under positive pressure, as resistance scales according to *R^-^*^4^ where *R* is vessel radius.

After normalizing capillary widths to the average value at the lowest magnitude pressure, we found that positive-pressure increases affected capillaries more so than negative-pressure increases (Fig. 4C(ii)), with notable reductions across the 9 cmH_2_O pressure regime. Significant differences between the pressure modes emerged at 12 cmH_2_O. This increased compression under positive pressure is likely due to the radially compressive, non-biologically relevant force exerted on the capillaries by the alveolar wall as positive transpulmonary pressure increases.

We find that alveolar diameters across all transpulmonary pressure overlap when compared between both pressure modes (Fig. 4D(i)). Upon normalizing the alveolar diameters to their values at 3 cmH_2_O, we observe alveoli under both conditions expand similarly regardless of pressure type (Fig. 4D(ii)). This overlap suggests that the respiratory function of the lung is conserved under both positive- and negative-pressure ventilation. Rather, it is likely the differences in radial vascular distension between the two pressure types that may play a stronger role in the physiological effects that arise between pressure modes. These findings highlight the distinct physiological effects of positive and negative pressure on capillary and alveolar behavior, demonstrating that negative-pressure ventilation may offer advantages in reducing capillary compression and improving blood flow, which could have important implications for lung function and therapeutic approaches.

## Discussion

In this study, we investigated how age-related remodeling of the mechanical properties of the lung alters the micromechanical behaviors of the lung in response to pulmonary pressure at the single alveolus and capillary scales. Importantly, by utilizing the crystal ribcage, we recapitulate the native environment of the in-situ lung tissue across the extremes of the murine lifespan and preserve the functional ability of the tissue to respond dynamically to changes in pressures. We make the following key observations, namely (i) the lumens in postnatal mice are stiffer than those in their young adult counterparts (ii) stiffer alveolar walls in postnatal mice undergo significantly less expansion upon exertion of positive transpulmonary pressure, (iii) aged lungs are stiffer than young adult lungs at increasing transpulmonary pressure, (iv) lumens across the lifespan compress similarly in response to administration of positive transpulmonary pressure, and (v) administration of positive transpulmonary pressure significantly reduces capillary diameter compared to administration of negative transpulmonary pressure.

As any variations in vascular distension would affect transport of oxygen, immune cells, and other substances to and from the lung, these age-related changes would have major impacts on the respiratory system, immune system, and overall health of the mouse. Yet, the functional implication of these age-related structural and mechanical changes has remained poorly understood. Previous studies have suggested that changes in function and micromechanics that occur with aging are best described by non-linear trends^1,2^, a theory that agrees with the varied age-related distension trends evident in our data: particularly, how postnatal lumens exhibit an increased Young’s modulus compared to young adult and aged counterparts, while postnatal alveoli undergo restricted expansion and smaller relative changes in diameter in response to transpulmonary pressure changes, compared to the more compliant young adult and aged alveoli.

This substantial difference in alveolar distension is potentially related to the elevated rate of alveolarization, or the genesis of new alveoli, that occurs during the postnatal phase^39^ and coincides with elevated cell migration^40^. The increased stiffness observed within alveolar walls may hold cues for epithelial migration and the subsequent formation of new alveoli. Formation of new alveoli continues later in life^41,42^, yet the decreased alveolar stiffness and high degree of overlap with adult tissue shown by our data suggests that if such a state reoccurs it is likely not informed by local stiffness. Rather, these alveoli may enlarge from their younger states in an attempt to retain normal gas exchange as age-related increases in ECM stiffness hidden at the parenchymal level hinder normal functionality^6^.

Our work here further adds to the ongoing discussion of positive- versus negative-pressure ventilation by examining the effect of positive and negative pressure on alveolar and capillary distension. While it is widely accepted that positive-pressure ventilation has deleterious patient effects compared to negative-pressure ventilation including brain damage^43^, ventilator induced lung injury^27–29^, and impaired renal function^44,45^, isolating the causes of these negative side effects has proven difficult. In ex vivo and in situ experiments, select studies suggest better oxygenation occurs in negative pressure than in positive pressure^33,35^, with CT imaging suggesting negatively ventilated lungs have more normally aerated tissue and less alveolar collapse (i.e., atelectasis)^35^. Other studies argue that the two forms have no effective difference in oxygenation or lung volume^23^ when pressure administration profiles are matched. Still, studies isolated to just the lungs show negative pressure produces better pulmonary blood flow^35^, as under positive pressure the pressure at the right atrium pressure is increased while cardiac output is reduced by 10-36%^30,46^. This finding aligns with our data, where capillary diameter decreases by approximately 15% across a positive transpulmonary increase of 9 cmH_2_O, a capillary change that corresponds to an almost doubling of resistance to the fluid flow, according to Poiseuille’s Law, while capillaries under negative transpulmonary pressure only undergo a 3% reduction across the 9 cmH_2_O transpulmonary pressure increase.

While our method has several advantages over traditional methods for exploring lung tissue mechanical properties, the complicated nature of the cardiopulmonary system currently presents some limits to the complexity of the model we can build. Namely, the measurements in our system are taken at quasistatic transpulmonary and vascular pressures. Future work should utilize dynamic circulation-respiration coupling methods to improve the complexity of the model and depth of data acquired: for example, quantifying vascular fluid flow in response to pressure changes. Additionally, future work can explore the effect of this micromechanical coupling on the immune system: for example, how positive- versus negative-pressure conditions affect pulmonary lymphatic flow or the trafficking of immune cells^24,47,48^, and how these relationships and dynamic readouts may change with age. Further exploration of pulmonary distension, its relationship to ventilation pressure, and its impact on immune response could have far-reaching implications, particularly given the effect of positive pressure ventilation and age-related mechanical changes observed in the brain^43,49–51^. Thus, our results here are a promising framework to continue to probe the mechanistic basis of the functional changes that occur in the lung with age.

## Materials and Methods

### Animal use ethics

All experiments adhered to ethical standards and guidelines under protocols approved by the Boston University Institutional Animal Care and Use Committee. Mice were housed and bred under pathogen-free environments at the Boston University Animal Science Center with appropriate ambient temperature, humidity, and light-dark conditions maintained.

### Mice

We used 6-7 day old (postnatal, unsexed), 12-18 week old (young adult, female), and 20+ month old (aged, female) C57BL/6 mice for the experimental procedures. A breeding pair of transgenic B6.129(Cg)-Gt(ROSA)26Sortm4(ACTB-tdTomato,-EGFP)Luo/J (JAX, 007676, Jackson Labs) with a transgenic tdTomato membrane signal was purchased to start a colony and was the source of all postnatal and young adult mice for the experiments. Aged wild type C57BL/6 mice were acquired from the National Institute on Aging (NIA) at the National Institutes of Health (NIH).

### Crystal ribcage design and fabrication

The crystal ribcage is a transparent biocompatible, age- and strain-matched physiological environment for imaging the ex vivo lung^8^. Detailed fabrication of the ribcage has been previously described^8^ and is summarized here: previously recorded microcomputed tomography scans of the mouse chest (C57BL/6 scans were obtained courtesy of the Hoffman group at the University of Iowa^52^) were used to define a 3D model of the chest cavity which was printed using a stereolithography printer (Form3, FormLabs). The printed model was polished to remove printing artifacts and then used as a positive mold to thermoform the crystal ribcage over. The base model was made age-specific by scaling the geometry by mouse lung volume variations with age^2^. Instead of plasma treating the crystal ribcage to engineer surface hydrophilicity as previously cited^8^, the internal surface was dip coated in iSurGlide Plus (iSurTech) following the manufacturer’s instructions to obtain a permanent, lubricious, and optically transparent hydrophilic coating on the crystal ribcage. Briefly, the outer surface of the ribcage was masked with a two-part silicone material (MoldStar 31T, Smooth-On), the masked ribcage surface was thoroughly cleaned with isopropanol and lint free Kimtech wipes (Kimberly-Clark Professional), the ribcage was dipped into the iSurGlide solution and allowed to dwell for 30 s before removing at a controlled rate of 3 mm/s, and the coated ribcages were allowed to degas and dry in a chemical fume hood for 5 minutes. The silicone mask was carefully removed before the coated ribcage was placed on a rotary stage 3 inches away from an 8W 302nm UV-B light source to cure for 15 minutes. The final coated ribcage is placed in a phosphate buffered solution to stay hydrated and maintain tonicity.

### Lung Preparation

Mice were anesthetized with a ketamine/xylazine cocktail and were ventilated (RoVent, Kent Scientific Corporation) through a tracheal cannula to prevent lung collapse before sacrificing the mouse. Lungs were then perfused with the perfusate medium (RPMI with 10% FBS, Gibco) supplemented with Evans Blue vascular dye (0.5 mg/ml, Fisher Scientific) through two cannulas inserted into the pulmonary artery and left atrium, with the pulmonary artery cannula attached to an external 10 ml reservoir of media. Evans Blue marks the vasculature and additionally ensures that the alveoli that are imaged are air-filled and not edematous.

After perfusion was established, the lung-heart block was removed from the chest cavity and placed into the crystal ribcage for microscopy. Perfusate was allowed to circulate at approximately 5-10 cmH_2_O of pressure driven flow for 5-10 minutes to label the lung’s interior vasculature before inflow and outflow were sealed at 0 cmH_2_O of hydrostatic vascular pressure. Afterwards, active ventilation was replaced with a positive transpulmonary pressure applied via external water column. Vascular pressure was controlled by varying the height of the external reservoir. Both transpulmonary and vascular pressures were confirmed manually and through pressure sensors.

For all negative pressure experiments, the lung preparation followed the same steps as described above. After the lung is placed in the crystal ribcage, the base of the ribcage was sealed using a silicone elastomer (Ecoflex 00-35 FAST, Smooth-On). Any gaps in the crystal ribcage or trachea opening were also sealed. Negative pressure was then applied via a syringe attached to the base of the ribcage and the trachea was then exposed to atmospheric pressure. Air pressure sensors confirmed the magnitude of negative pressure applied to the crystal ribcage.

### Lung Microscopy

Ex vivo lungs inside the crystal ribcage were imaged under quasi-static inflation and perfusion conditions using an upright spinning disc confocal microscope (Nikon CSU X1) with a 10x objective. Signal from Evans Blue was captured using a 640nm laser coupled with a Cy5 far red emission filter, with laser power set to 10-15% and exposure at 50-100ms. Signal from the tdTomato expressing transgenic reporter mice was collected with a 561nm laser coupled with TRITC (tetramethylrhodamine) filter, with laser power set to 0.5% and exposure between 20- 50ms. Imaging was focused to the base of the upper right lobe of the lung for consistency across experiments and was comprised of Z stacks using the objective’s optimal Z spacing and XY resolution for voxels of 0.65 x 0.65 x 2.5 µm^3^. Additional widefield images were obtained at the benchtop using an upright Nikon stereomicroscope. Microscopy data were processed and visualized using FIJI, custom written Python codes, and MATLAB R2024b as outlined below.

### Quasi-static vascular distention measurements

For vascular distension measurements, ex vivo lungs inside the crystal ribcage were kept at a constant transpulmonary pressure of 7 cmH_2_O via trachea cannula attached to external water column. Vascular pressure was varied by raising the perfusion reservoir to create height differences between the surface of the liquid and the approximate location of the heart: this height difference was measured manually and confirmed with inflow pressure sensor. The reservoir height was adjusted while tubing from both cannulas into and out of the heart was open, allowing free media flow through the system, before the cannula inserted into the left atrium was clamped closed to stop outflow of perfusate. Images were taken once the vascular pressure measured at both cannulas equilibrated. These pressure changes are of a quasistatic nature that may not fully capture the dynamic pressure changes of the in vivo circulation- respiration, yet this remains the best approach for directly estimating the pressure within the capillaries.

### Quasi-static alveolar distention measurements

Ex vivo lungs inside the crystal ribcage were kept at a constant vascular pressure of 0 cmH_2_O while transpulmonary pressure was varied by either positive or negative pressure. For positive-pressure experiments, transpulmonary pressure was varied using a water column to control pressure at the trachea while the bottom of the crystal ribcage was open to atmospheric pressure. For negative pressure experiments, the base of the rib cage was sealed, and pressure was controlled via a syringe attached to the base of the ribcage while the trachea was open to atmospheric pressure (0 cmH_2_O). The lung pressure was able to be switched from positive to negative pressure without collapsing the lung. Transpulmonary pressure values were verified using sensors attached to either the trachea (positive pressure) or the base of the ribcage (negative pressure). These quasistatic air pressure changes may not fully capture the dynamics of pressure change in circulation-respiration, yet they remain the best approach for directly estimating the pressure within the alveoli.

### Image segmentation, matching, and measurements

Full description of image processing may be found in Supplemental Information. A brief description follows here: microscopy images of lung tissue were converted to 8-bit summed z projections using FIJI before being analyzed using a custom Python-based image processing pipeline. Segmentation was performed using a fine-tuned Segment Anything Model (SAM)^36^, with preprocessing steps including intensity normalization and contrast enhancement. We fine-tuned SAM to then automatically detect and outline the perimeter of each alveolus. We then complete affine image registration between images at each pressure level, using manual selection of at least 3 corresponding landmarks per image. Finally, SAM was used to identify and measure the diameters of matched alveoli and septa.

Alveolar measurements were conducted through centroid-based correlation analysis of the segmented regions. For each identified alveolus, the diameter was calculated from the segmented area. Spatial relationships between adjacent alveoli were quantified using Delaunay triangulation of centroid coordinates. Distances across each lumen were measured by analyzing inter-alveolar regions, with 300 sampling points along each potential lumen edge boundary. All measurements underwent manual validation through a custom graphical interface and were exported for statistical analysis.

The analysis pipeline was implemented in Python 3.8, utilizing *OpenCV*, *scikit-image*, and *segment-anything* libraries. All image processing parameters were kept consistent across analyzed samples to ensure measurement comparability.

### Mathematical modeling of the mechanics of pulmonary capillaries

We model the solid phase of each pulmonary capillary as an infinitely long, thick-walled, linearly elastic, hollow cylinder. Epithelial cells and the surrounding basement membrane together comprise the walls of the pulmonary capillaries, whose stiffness arises primarily from a reticulated network of collagen fibrils within the basement membrane. Directional variations in the geometry of this collagen network may produce differences in the capillary’s circumferential and longitudinal compliance. Consequently, we assume that the material is transversely isotropic, with the plane of isotropy corresponding to the plane perpendicular to the symmetry axis of the cylinder. The general constitutive equation for a linearly elastic material is given by equation (1) where ε is the Cauchy strain tensor, *C* is the elastic compliance tensor, and σ is the elastic stress tensor.

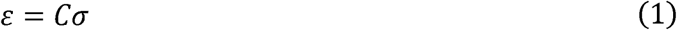

We assume that the cylinder is subject to uniform pressures of different magnitudes, *P_a_* and *P_b_*, on its inner and outer surfaces, as well as to uniform axial strain, E_ZZ_, along its length. Changes in arterial pressure correspond to changes in *P_a_*, while changes in alveolar pressure correspond to changes in *P_b_* . Due to the cylindrical symmetry of the geometry and the loading conditions, we adopt a cylindrical coordinate system whose z-axis is coincident with the symmetry axis of the cylinder. In this coordinate system, the constitutive equation simplifies considerably to equation (2).

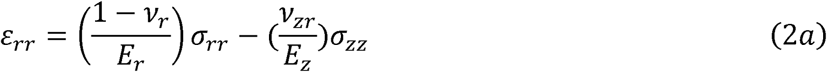

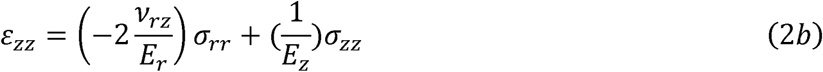

Given the constitutive equation, boundary conditions, and chosen system of coordinates, the equilibrium condition implies the radial displacements provided by equation (3), where a and b are the inner and outer radius in the reference configuration, *v_r_* is the Poisson ratio in the plane of isotropy, *v_rZ_* is the Poisson ratio characterizing axial strain due to radial stress, *v_Zr_* is the Poisson ratio characterizing radial strain due to axial stress, *E_r_* is the radial stiffness, and *E_Z_* is the axial stiffness.

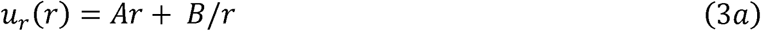

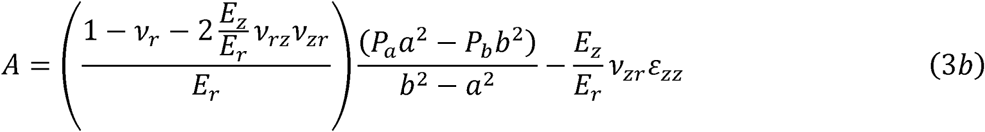

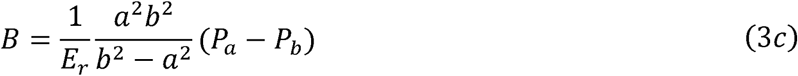

For simplicity, we set each Poisson ratio to zero, which decouples the radial strain from the axial stiffness and the axial strain from the radial stiffness. This enables us to determine the axial stiffness from the vascular-distension experiments, where the magnitude of the radial strain is much larger than that of the axial strain, and the radial stiffness from the alveolar-distension experiments. When the Poisson ratio is zero, the radial-displacement equation (3a) simplifies to equation (4).

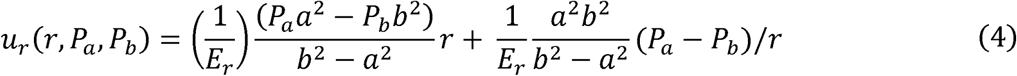

During our vascular-distension experiments, we increase the arterial pressure *P_a_* from *P_a,1_* to *P_a,2_*, while maintaining constant alveolar pressure *P_b_*. Thus, the radial displacement of the inner surface changes by *Δu_r_* = *u_r_* (*a, P_a,2_*, *P_b_* )- *u_r_*(*a, P_a,1_*, *P_b_*), which simplifies as shown in equation (5).

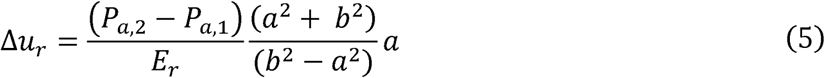

We solve this equation for the radial stiffness, producing equation (6).

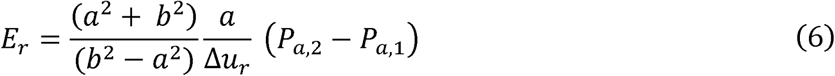

When the Poisson ratio is zero, we can solve for the axial stiffness from the axial strain equation (2b), yielding equation (7).

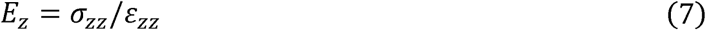

To determine the axial stiffness from this equation, we can measure the axial strain, but we must estimate the axial stress, which is not directly measurable. During our alveolar-distension experiments, we increase the alveolar pressure *P_b_* from *P_b,1_* to *P_b,2_*, while maintaining constant vascular pressure *P_a_*. Following Mead^53^, we assume that under quasi-static conditions, the average change in stress on the pleural surface equals the change in transpulmonary pressure, *P_b,2_* - *P_b,1_*. Consider a thin region of the lung containing the pleural surface. To maintain equilibrium, if the capillary wall is the primary load-bearing element, then the net tensile force within the capillary wall must precisely balance the net force produced by the transpulmonary pressure across the pleural surface, leading to equation (8).

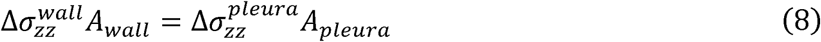

The axial stress within the capillary wall is then given by equation (9), where *A_pleura_* is the area of a region of the pleural surface and *A_wall_* is the area of the capillary wall over the same region.

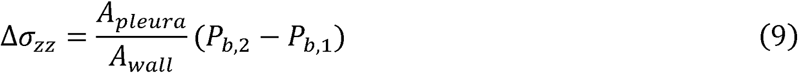

We approximate this area ratio, 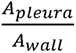, as follows. First, we expand the ratio as shown in equation (10), because the ratios 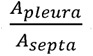 and 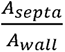 are simple to approximate.

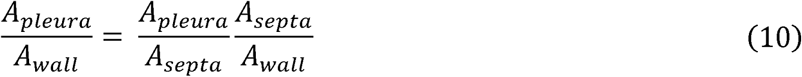

If we assume that the septum geometry resembles a regular hexagonal lattice, then it can be shown that the ratio 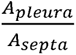 is approximately given by equation (11), the derivation of which is explained further in Supplemental Information, where L is the side length of each hexagon and *R* is half the thickness of each hexagon’s side.

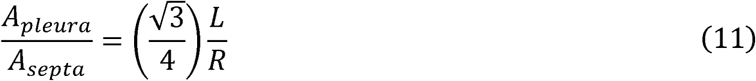

Meanwhile, to approximate the ratio 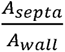, we assume that the septum is entirely composed of hollow, thick-walled, cylindrical capillaries. Thus, this ratio is approximated as the area of the disk with radius *b*, i.e. *πb*^2^, divided by the area of the annulus with inner radius a and outer radius *b*, i.e. *π(b*^2^ - *a*^2^). This yields equation (12).

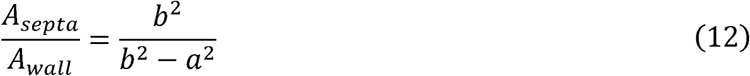

By combining equations (9)-(12), and setting *R=b,* we recover the approximate stress within the capillary wall, given by equation (13).

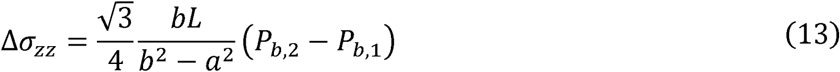

By substituting equation (13) into equation (7), we recover our final expression for the axial stiffness, as shown in equation (14).

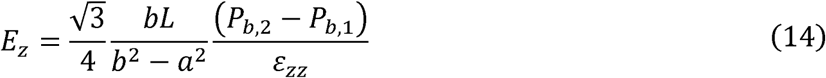

## Statistics and reproducibility

All experiments are performed in at least four separate mice to generate biologically independent samples. Exact sample sizes are presented in the captions of each figure where the data is presented; data is presented as mean + standard error of the mean (s.e.m.). To obtain normalized measurements, the average value at each pressure was calculated per mouse, and then divided by the average value at the lowest magnitude pressure measured. Data from each individual mouse was then averaged together to obtain the reported values for each experiment. All statistics presented here were calculated as a two-tailed Student’s t-test to determine P values between groups of data, with differences considered significant when P < 0.05 and trends when P < 0.10. Exact P values comparing postnatal to young adult, and young adult to aged groups are shown in plots. Statistical analyses were performed in MATLAB R2024b.

## Data Availability Statement

The data used in this study is available by the corresponding author upon reasonable request.

## Code Availability Statement

The code and training data used in this study is available by the corresponding author upon reasonable request.

## Acknowledgements

We acknowledge the following research support: NIH R21EB031332 and DP2HL168562, Sloan Research Fellowship, Beckman Young Investigator Award, NSF CAREER, Boston University CMTM and Dean’s Catalyst Awards, and the American Cancer Society Institutional Fund at Boston University to H.T.N; Kilachand Fund to H.T.N.; NIH F30HL168952 to L.S.; NSF GRFP to G.N.G.; NIH S10OD024993 to Boston University BME Department. This research was supported in part by the National Institutes of Health training grant at Boston University, T32 EB006359. The content is solely the responsibility of the authors and does not necessarily represent the official views of the National Institutes of Health.

## Author Contributions

K.R. and L.C. designed and performed the experiments, performed the analysis, and wrote the manuscript with equal contribution. R.L. designed, implemented, and analyzed the axial and radial stiffness modeling, and additionally processed experimental analysis. A.K. designed and optimized the computational analysis pipeline for image segmentation, registration, and measurement. L.S. prepared the postnatal samples for experiment and imaging. W.W. manufactured the crystal ribcages. G.G. contributed to negative pressure and vascular pressure experimental designs. R.B. supported crystal ribcage sensors & software. A.B. manufactured postnatal crystal ribcages. B.S. contributed to analysis discussions and guidance. H.T.N. designed the study, supported and interpreted the analysis, and revised the manuscript.

## References

1 Huang, K., Rabold, R., Schofield, B., Mitzner, W. & Tankersley. Age-dependent changes of airway and lung parenchyma in C57BL/6J mice. Journal of Applied Physiology 102, 200–206 (2007). 10.1152/japplphysiol.00400.2006

2 Schulte, H., M ü hlfeld, C. & Brandenberger, C. Age-Related Structural and Functional Changes in the Mouse Lung. Frontiers in Physiology 10 (2019).

3 Janssens, J. P., Pache, J. C. & Nicod, L. P. Physiological changes in respiratory function associated with ageing. European Respiratory Journal 13, 197–205 (1999). 10.1034/j.1399-3003.1999.13a36.x

4 Sicard, D., Haak, A., Choi, K. M., Craig, A., Fredenburch, L. & Tschumperlin, D. Aging and anatomical variations in lung tissue stiffness. American Journal of Physiology-Lung Cellular and Molecular Physiology 314, L909–L1025 (2018).

5 Subramanian, K., Kumar, H. & Tawhai, M. H. Evidence for age-dependent air-space enlargement contributing to loss of lung tissue elastic recoil pressure and increased shear modulus in older age. Journal of Applied Physiology 123, 79–87 (2017). 10.1152/japplphysiol.00208.2016

6 Suki, B. & Bartolák-Suki, E. in Mechanical Properties of Aging Soft Tissues (eds B. Derby & R. Akhtar) (Springer, Cham., 2015).

7 Schneider, J. L., Rowe, J. H., Garcia-de-Alba, C., Kim, C. F., Sharpe, A. H. & Haigis, M. C. The aging lung: physiology, disease, and immunity. Cell 184, 1990–2019 (2021). 10.1016/j.cell.2021.03.005

8 Banerji, R., Grifno, G. N., Shi, L., Smolen, D., LeBourdais, R., Muhvich, J., Eberman, C., Hiller, B., Lee, J., Regan, K., Zheng, S., Zhang, S. S., Jiang, J., Breda, J. C., Pihl, R., Traber, K., Mazzilli, S., Ligresti, G., Mizgerd, J. P., Suki, B. & Nia, H. T. Crystal ribcage: a platform for probing real-time lung function at cellular resolution in health and disease. Nature Methods (2023). 10.1038/s41592-023-02004-9

9 Suki, B. & Bates, J. H. T. Lung tissue mechanics as an emergent phenomenon. Journal of Applied Physiology 110, 1111–1118 (2011). 10.1152/japplpysiol.01244.2010

10 Hirota, N. & Martin, J. G. Mechanisms of airway remodeling. Chest 144, 1026–1032 (2013). 10.1378/chest.12-3073

11 Malhotra, R., Dhakal, B., Eisman, A., Pappagianopoulos, P., Dress, A., Weiner, R., Baggish, A., Semigran, M. & Lewis, G. Pulmonary Vascular Distensibility Predicts Pulmonary Hypertension Severity, Exercise Capacity, and Survival in Heart Failure. Circulation: Heart Failure 9 (2017). 10.1161/CIRCHEARTFAILURE.115.003011

12 Roan, E. & Waters, C. What do we know about mechanical strain in lung alveoli? American Journal of Physiology- Lung Cellular and Molecular Physiology 301, L625–L828 (2011).

13 Bachofen, H. & Sch ürch, S. Alveolar surface forces and lung architecture. Comparative Biochemistry and Physiology Part A: Molecular & Integrative Physiology 129, 183–193 (2001).

14 Peták, F., Janosi, T., Myers, C., Fontao, F. & Habre, W. Impact of elevated pulmonary blood flow and capillary pressure on lungresponsiveness. Journal of Applied Physiology 107, 639–1003 (2009).

15 Waters, C., Sporn, P., Liu, M. & Jeffrey, F. Cellular biomechanics in the lung. American Journal of Physiology-Lung Cellular and Molecular Physiology 283, L503–L663 (2002).

16 Wasserman, K., Zhang, Y.-Y., Gitt, A., Belardinelli, R., Koike, A., Lubarsky, L. & Agostoni, P. G. Lung Function and Exercise Gas Exchange in Chronic Heart Failure. Circulation 96, 2221–2227 (1997).

17 Ryans, J. M., Fujioka, H. & Gaver, D. P. Microscale to mesoscale analysis of parenchymal tethering: the effect of heterogenous alveolar pressures on the pulmonary mechanics of compliant airways. Journal of Applied Physiology 126, 1204–1213 (2019).

18 Zhang, S., Grifno, G. N., Passaro, R., Regan, K., Zheng, S., Hadzipasic, M., Banerji, R., O’Connor, L., Chu, V., Kim, S. Y., Yang, J., Shi, L., Karrobi, K., Roblyer, D., Grinstaff, M. W. & Nia, H. T. Intravital measurements of solid stresses in tumours reveal length-scale and microenvironmentally dependent force transmission. Nature Biomedical Engineering (2023). 10.1038/s41551-023-01080-8

19 Ekpenyong, A. E., Toepfner, N., Fiddler, C., Herbig, M., Li, W., Cojoc, G., Summers, C., CGuck, J. & Chilvers, E. R. Mechanical deformation induces depolarization of neutrophils. Science Advances 3 (2017). 10.1126/sciadv.1602536

20 Solis, A. G., Bielecki, P., Steach, H. R., Sharma, L., Harman, C. C. D., Yun, S., de Zoete, M. R., Warnock, J. N., To, S. D. F., York, A. G., Mack, M., Schwartz, M. A., Dela Cruz, C. S., Palm, N. W., Jackson, R. & Flavell, R. A. Mechanosensation of cyclical force by PIEZO1 is essential for innate immunity. Nature 573, 69–74 (2019). 10.1038/s41586-019-1485-8

21 Aykut, B., Chen, R., Kim, J. I., Wu, D., Shadaloey, S. A. A., Abengozar, R., Preiss, P., Saxena, A., Pushalkar, S., Leinwand, J., Diskin, B., Wang, W., Werba, G., Berman, M., Lee, S. K. B., Khodadadi-Jamayran, A., Saxena, D., Coetzee, W. A. & Miller, G. Targeting Piezo1 unleashes innate immunity against cancer and infectious disease. Sci Immunol 5 (2020). 10.1126/sciimmunol.abb5168

22 Lefrançais, E., Ortiz-Muñoz, G., Caudrillier, A., Mallavia, B., Liu, F., Sayah, D. M., Thornton, E. E., Headley, M. B., David, T., Coughlin, S. R., Krummel, M. F., Leavitt, A. D., Passegué, E. & Looney, M. R. The lung is a site of platelet biogenesis and a reservoir for haematopoietic progenitors. Nature 544, 105–109 (2017). 10.1038/nature21706

23 Engelberts, D., Malhotra, A., Butler, J. P., Topulos, G. P., Loring, S. H. & Kavanagh, B. P. Relative effects of negative versus positive pressure ventilation depend on applied conditions. Intensive Care Medicine 38, 879–885 (2012). 10.1007/s00134-012-2512-5

24 Burrowes, K. S., Tawhai, M. H. & Hunter, P. J. Modeling RBC and neutrophil distribution through an anatomically based pulmonary capillary network. Annals of Biomedical Engineering 32, 585–595 (2004). 10.1023/b:abme..0000019178.95185.ad

25 Kim, J., Heise, R. L., Reynolds, A. M. & Pidaparti, R. Quantification of Age-Related Lung Tissue Mechanics under Mechanical Ventilation. Medical Sciences 5, 21 (2017). 10.3390/medsci5040021

26 Aghasafari, P., Heise, R., Reynolds, A. & Pidaparti, R. Aging Effects on Alveolar Sacs Under Mechanical Ventilation. The Journals of Gerontology Series A 74 (2018).

27 Gattinoni, L., Tonetti, T. & Quintel, M. Regional physiology of ARDS. Critical Care 21 (2017).

28 Slutsky, A. S., M.D. & Ranieri, V. M., M.D. Ventilator-Induced Lung Injury. New England Journal of Medicine 369, 2126–2136 (2013).

29 Minucci, S., Heise, R. L., Valentine, M. S., Kamga Gninzeko, F. J. & Reynolds, A. M. Mathematical modeling of ventilator-induced lung inflammation. Journal of Theoretical Biology 526 (2021). 10.1016/j.jtbi.2021.110738

30 Corp, A., Thomas, C. & Adlam, M. The cardiovascular effects of positive pressure ventilation. BJA Education 21, 202–209 (2021). 10.1016/j.bjae.2021.01.002

31 Frank, J. & Matthay, M. Science review: Mechanisms of ventilator-induced injury. Critical Care 7, 233–241 (2003).

32 Kelly, G., Faraj, R., Zhang, Y., Maltepe, E., Fineman, J., Black, S. & Wang, T. Pulmonary Endothelial Mechanical Sensing and Signaling, a Story of Focal Adhesions and Integrins in Ventilator Induced Lung Injury. Frontiers in Physiology 10 (2019).

33 Sattari, S. Positive- and Negative-Pressure Ventilation Characterized by Localand Global Pulmonary Mechanics. American Journal of Respiratory and Critical Care MedicineVolume 207, Issue 5 207 (2023).

34 Quiros, K. A. M., Nelson, T. M., Ulu, A., Dominguez, E. C., Biddle, T. A., Lo, D. D., Nordgren, T. M. & Eskandari, M. A Comparative Study of Ex[1]Vivo Murine Pulmonary Mechanics Under Positive[1] and Negative[1]Pressure Ventilation. Annals of Biomedical Engineering 52, 342–354 (2024).

35 Grasso, F., Engelberts, D., Helm, E., Frndova, H., Jarvis, S., Talakoub, O., McKerlie, C., Babyn, P., Post, M. & Kavanagh, B. P. Negative-Pressure Ventilation: Better oxygenation and less injury. American Journal of Respiratory and Critical Care Medicine 177 (2007). 10.1164/rccm.200707-1004OC

36 Kirillov, A., Mintun, E., Ravi, N., Mao, H., Rolland, C., Gustafson, L., Xiao, T., Whitehead, S., Berg, A. C., Lo, W.-Y., Dollár, P. & Girshick, R. (arXiv:2304.02643, 2023).

37 Thibault, H. B., Kurtz, B., Raher, M. J., Shaik, R. S., Waxman, A., Derumeaux, G., Halpern, E. F., Bloch, K. D. & Scherrer-Crosbie, M. Noninvasive assessment of murine pulmonary arterial pressure: validation and application to models of pulmonary hypertension. Circulation: Cardiovascular Imaging 3, 157–163 (2010). 10.1161/circimaging.109.887109

38 Jawde, S. B., Takahashi, A., Bates, J. H. T. & Suki, B. An Analytical Model for Estimating Alveolar Wall Elastic Moduli From Lung Tissue Uniaxial Stress-Strain Curves. Frontiers in Physiology (2020). 10.3389/fphys.2020.00121

39 Mund, S. I., Stampanoni, M. & Schittny, J. C. Developmental alveolarization of the mouse lung. Developmetanl Dynamics 237, 2108–2116 (2008).

40 Akram, K. M., Yates, L. L., Mongey, R., Rothery, S., Gaboriau, D. C. A., Sanderson, J., Hind, M., Griffiths, M. & Dean, C. H. Live imaging of alveologenesis in precision-cut lung slices reveals dynamic epithelial cell behavior. Nature Communications 10 (2019). 10.1038/s41467-019-09067-3

41 Schittny, J. C. Development of the Lung. Cell and Tissue Research 367, 427–444 (2017). 10.1007/s00441-016-2545-0

42 Pozarska, A., Rodríguez-Castillo, J. A., Solalingue, D. E. S., Ntokou, A., Rath, P., Mižíková, I., Madurga, A., Mayer, K., Vadász, I., Herold, S., Ahlbrecht, K., Seeger, W. & Morty, R. E. Stereological monitoring of mouse lung alveolarization from the early postnatal period to adulthood. American Journal of Physiology-Lung Cellular and Molecular Physiology 312, L882–895 (2017). 10.1152/ajplung.00492.2016

43 Bassi, T. G., Rohrs, E. C., Fernandez, K. C., Ornowska, M., Nicholas, M., Gani, M., Evans, D. & Reynolds, S. C. Brain injury after 50 h of lung-protective mechanical ventilation in a preclinical model. Scientific Reports 11 (2021). 10.1038/s41598-021-84440-1

44 Hepokoski, M. L., Malhotra, A., Singh, P. & Crotty Alexander, L. E. Ventilator Induced Kidney Injury: Are novel biomarkers the key to prevention? Nephron 140, 90–93 (2018). 10.1159/000491557

45 van den Akker, J. P., Egal, M. & Groeneveld, A. J. Invasive mechanical ventilation as a risk factor for acute kidney injury in the critically ill: a systematic review and meta-analysis. Critical Care 17 (2013). 10.1186/cc12743

46 Bindslev, L., Hedenstrierna, G., Santesson, J., Gottlieb, I. & Carvallhas, A. Ventilation-Perfusion Distribution During Inhalation Anaesthesia. Acta Anaesthesiologica Scandinavica 25, 360–371 (1981). 10.1111/j.1399-6576.1981.tb01667.x

47 Li, H., Zhang, J., Padera, T. P., Baish, J. W. & Munn, L. L. Fluid dynamics and leukocyte transit in the lymphatic system. PNAS Nexus 3 (2024). 10.1093/pnasnexus/pgae195

48 Cleary, S. J., Qiu, L., Seo, Y., Baluk, P., Liu, D., Serwas, N. K., Taylor, C. A., Zhang, D., Cyster, J. G., McDonald, D. M., Krummel, M. F. & Looney, M. R. Intravital imaging of pulmonary lymphatics in inflammation and metastatic cancer. Journal of Experimental Medicine 222, e20241359 (2025). 10.1084/jem.20241359

49 Zheng, S., Banerji, R., LeBourdais, R., Zhang, S., DuBois, E., O’Shea, T. & Nia, H. Alteration of mechanical stresses in the murine brain by age and hemorrhagic stroke. PNAS Nexus 3 (2024). 10.1093/pnasnexus/pgae141

50 Lv, H., Kurt, M., Zeng, N., Ozkaya, E., Marcuz, F., Wu, L., Laksari, K., Camarillo, D. B., Pauly, K. B., Wang, Z. & Wintermark, M. MR elastography frequency-dependent and independent parameters demonstrate accelerated decrease of brain stiffness in elder subjects. European Radiology 30, 6614–6623 (2020).

51 Ozkaya, E., Fabris, G., Macruz, F., Suar, Z. M., Abderezaei, J., Su, B., Laksari, K., Wu, L., Camarillo, D. B., Pauly, K. B., Wintermark, M. & Kurt, M. Viscoelasticity of children and adolsecent brains throuhg MR elastography. Journal of the Mechanical Behavior of Biomedical Materials 115 (2021). 10.1016\j.jmbbm.2020.104229

52 Kizhakke Puliyakote, A. S., Valilescu, D. M., Newell Jr., J. D., Wang, G., Weibel, E. R. & Hoffman, E. A. Morphometric differences between central vs. surface acini in A/J mice using high-resolution micro-computed tomography. Journal of Applied Physiology 121, 115–122 (2016). 10.1152/japplphysiol.00317.2016

53 Mead, J., Takishima, T. & Leith, D. Stress distribution in lungs: a model of pulmonary elasticity. Jornal of Applied Physiology 28 (1970).

