## Supplemental Figures for "Micromechanics of lung capillaries across mouse lifespan and in positive- vs negative- pressure ventilation"

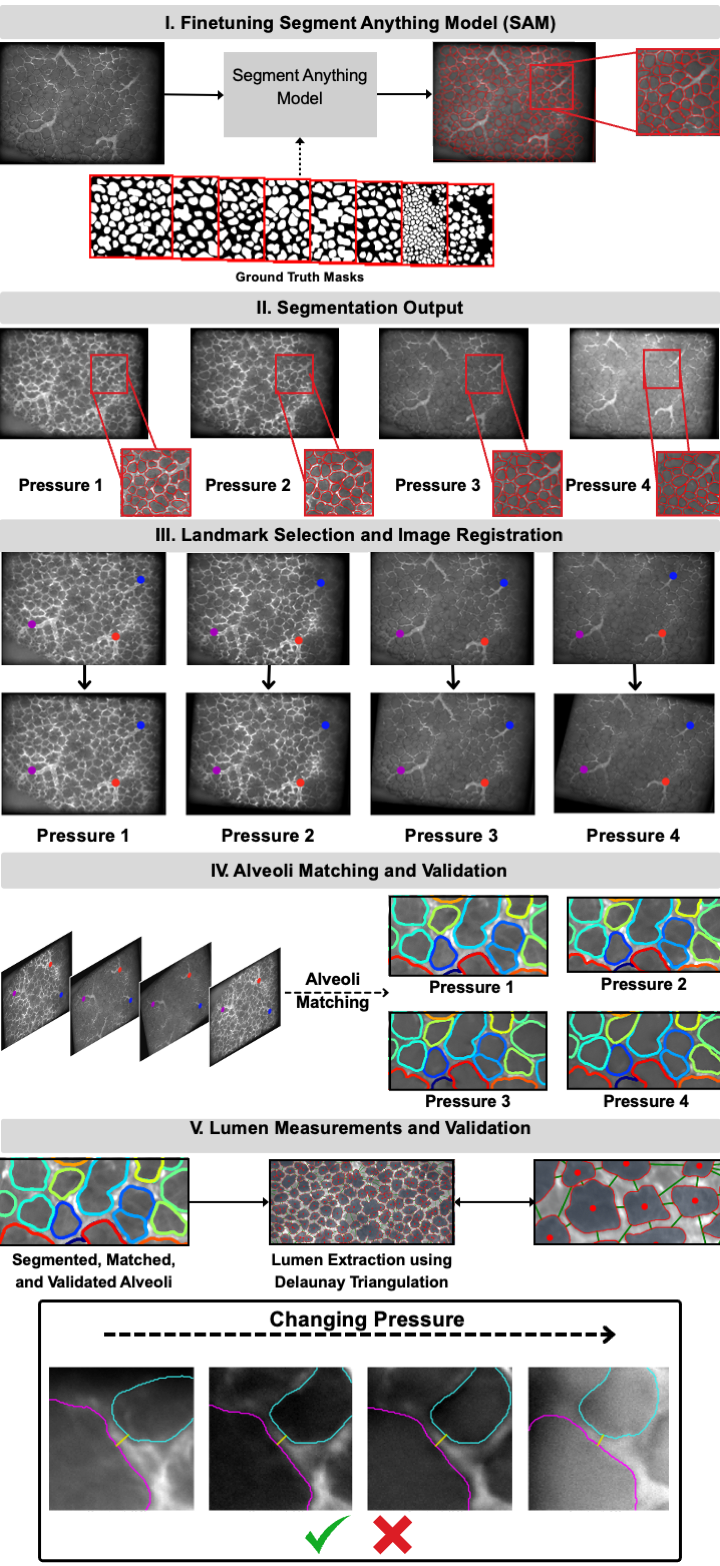

**Figure S1 | Pipeline for analyzing alveolar distention under varying pressure conditions using deep learning-based segmentation and registration.** The workflow consists of: (i) fine-tuning a Segment Anything Model (SAM) using ground truth, manually annotated masks of alveoli, (ii) automated segmentation of alveolar structures across group pressure conditions, visualized by red overlays on microscopy images, (iii) landmark-based registration using three user-selected anatomical markers (red, blue, purple) to align images across pressure states, (iv) centroid-based matching of individual alveoli across pressure conditions with user validation, and (v) extraction of alveolar lumen measurements via Delaunay triangulation from matched alveoli. The images in the bottom panel are shown to the user during the validation of the extracted lumen measurements. This semi-automated approach enables robust quantification of alveolar mechanics during respiration.

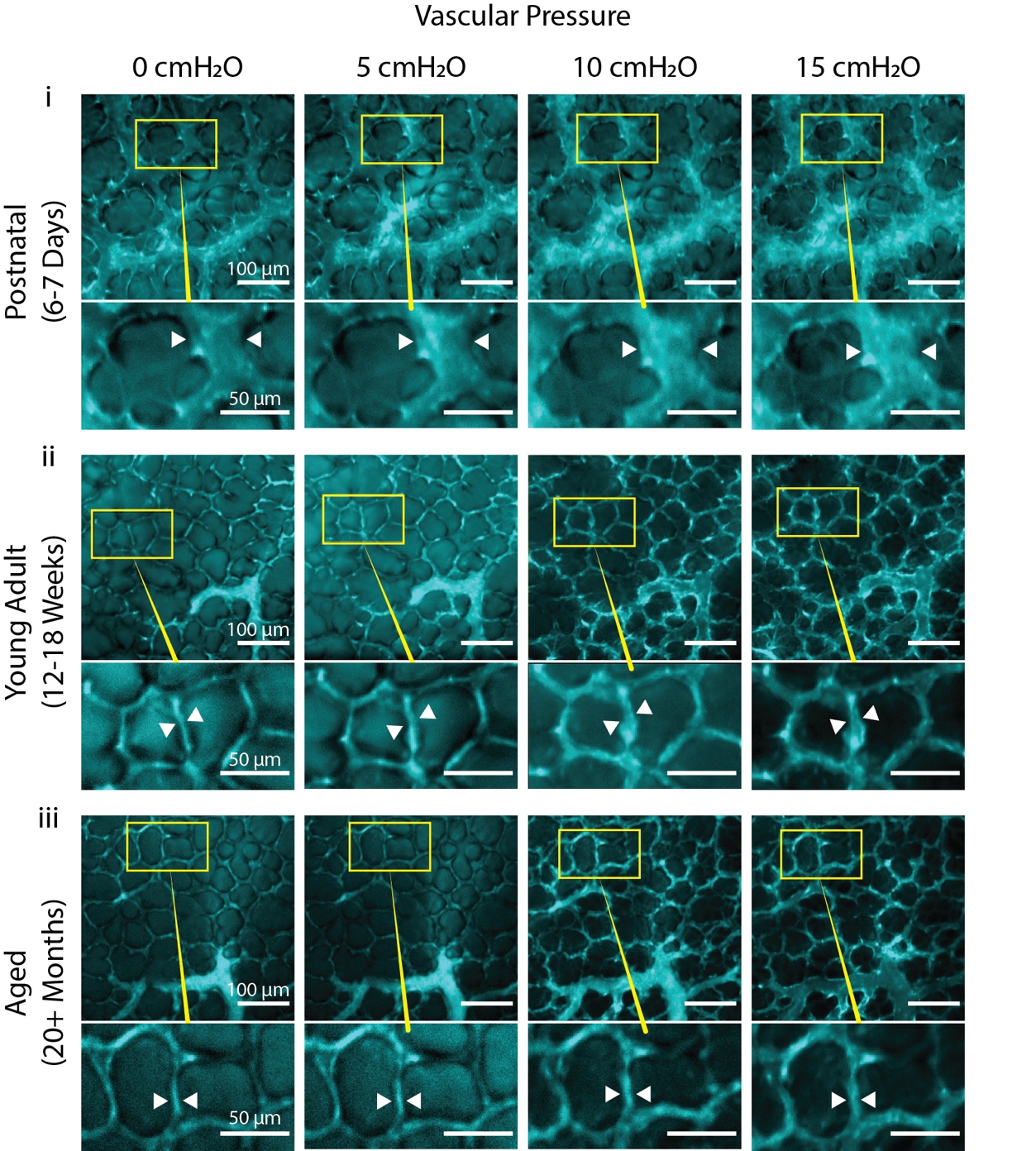

**Figure S2 | Aged lungs show smaller increase in lumen diameter as vascular pressure is increased across full dataset.** Confocal images capture changes in diameters of vessel lumens (white arrowheads) in response to vascular pressure changes. Lumens are labeled with Evans Blue dye via vascular perfusate. Shown are representative images of postnatal (i) young adult (ii) and aged (iii) vessels at vascular pressures of 0, 5, 10 and 15 cmH_2_O.

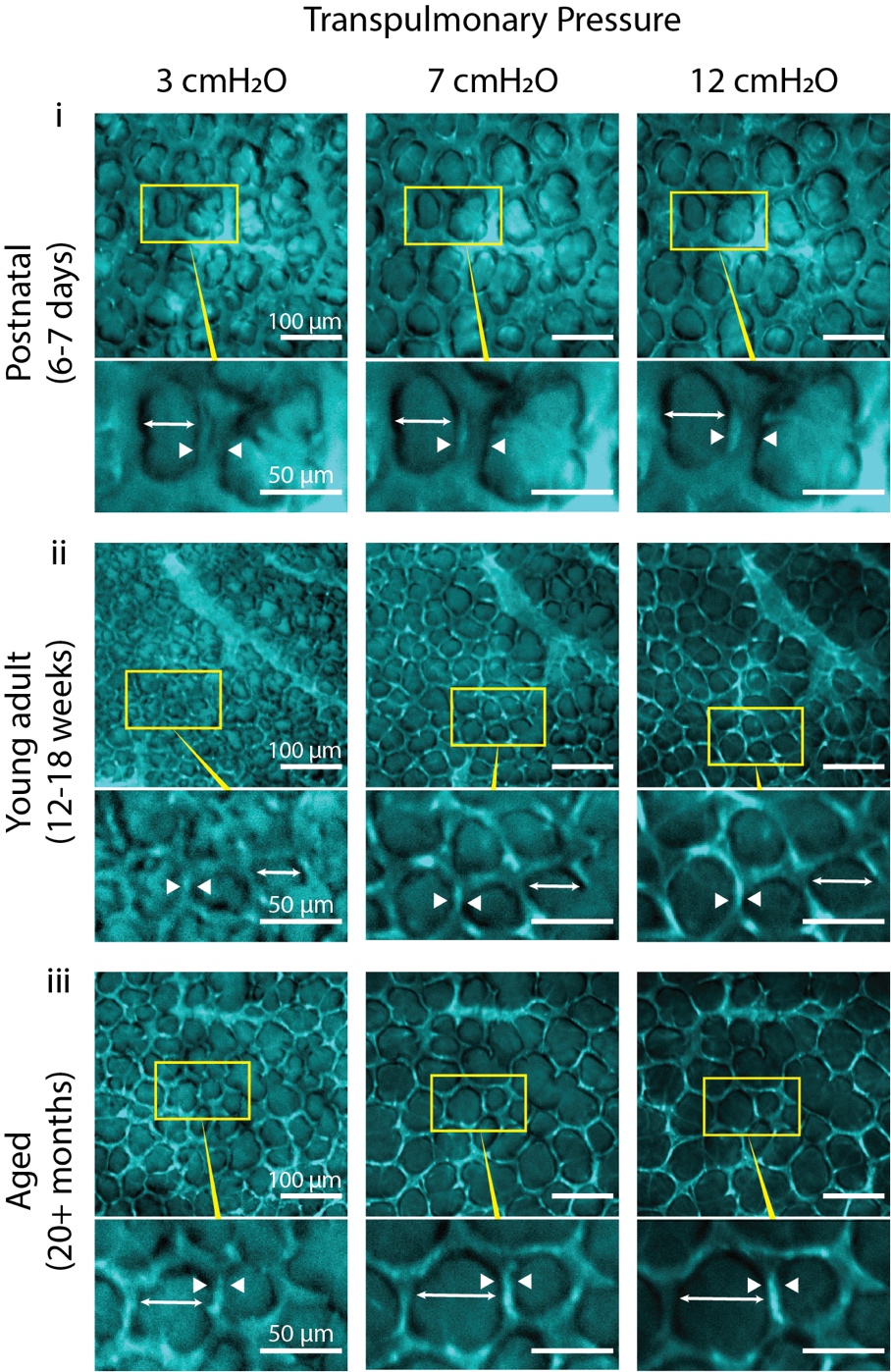

**Figure S3 | Postnatal alveoli show smaller diameter change in response to increasing transpulmonary pressure across the full dataset.** Confocal microscopy images of the vessels perfused with Evans Blue dye depict the alveolar level response to increases in positive transpulmonary pressure in postnatal (i), young (ii) and aged (iii) mice. At lower transpulmonary pressures, the diameter of the alveolus (white arrows) is lower than at higher, while the diameter of the lumen (white arrowheads) experiences the opposite effect.

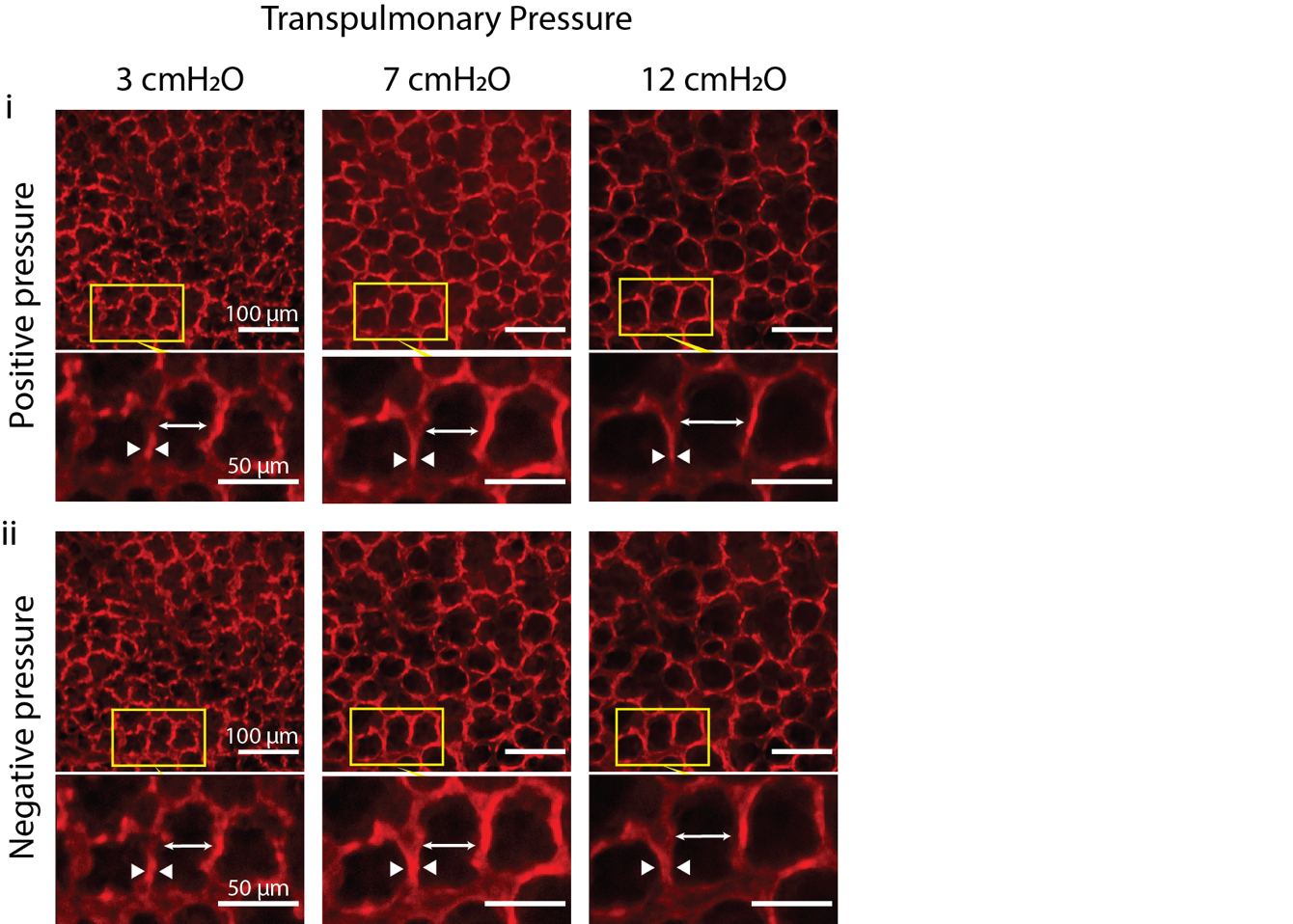

**Figure S4 | Capillaries of young mice show decreased change in diameter under negative pressure compared to positive across the full dataset.** Confocal microscopy images depict alveolar level response to increasing transpulmonary pressure under both positive- (i) and negative-pressure (ii) conditions. All images are taken at the same ROI within the same lung as the transpulmonary pressure is changed. At lower transpulmonary pressures, the diameter of the alveolus (white arrows) is lower than at higher, while the diameter of the lumen (white arrowheads) experiences the opposite effect.
