## Supplemental Information for "Micromechanics of lung capillaries across mouse lifespan and in positive- vs negative- pressure ventilation"

**Supplemental Information 1: Image Segmentation, matching, and measurements using AI-based segmentation algorithms**

Microscopy images of lung tissue were analyzed using a custom Python-based image processing pipeline. The analysis workflow consisted of three main stages: automated segmentation, image registration, and quantitative measurements of alveolar structures and septal distances.

Segmentation was performed using a fine-tuned Segment Anything Model (SAM) with a Vision Transformer (ViT-H) architecture^1^. Prior to segmentation, images underwent preprocessing which included intensity normalization and contrast enhancement using Contrast Limited Adaptive Histogram Equalization (CLAHE). The SAM model was configured with optimized parameters to ensure accurate detection of alveolar boundaries. The segmentation output was validated through visual inspection with contour overlays on the original images.

For comparative analysis across different tissue depths, affine image registration was implemented. Registration was performed using manually selected corresponding landmarks across all images. A minimum of three corresponding points were identified in each image set to compute the transformation. The transformation was optimized to align corresponding landmarks while maintaining structural integrity of the tissue features.

Alveolar measurements were conducted through centroid-based correlation analysis of the segmented regions. For each identified alveolus, the diameter was calculated assuming circular approximation, derived from the total area of the segmented region. The spatial relationships between adjacent alveoli were quantified using Delaunay triangulation of the centroid coordinates.

Septal distances were measured using an automated analysis of the inter-alveolar regions. The algorithm sampled 300 equally spaced points along each potential septal boundary, classifying regions as septal tissue based on the binary segmentation mask. The total septal length was calculated by summing the classified septal portions. To maintain measurement quality, all automatically detected septa underwent manual validation through a custom graphical interface that displayed both close-up and wide-field views of each septum.

All measurements were exported to CSV format for subsequent statistical analysis. The analysis pipeline was implemented in Python 3.8, utilizing OpenCV for image processing, scikit-image for morphological operations, and specialized libraries including segment-anything for deep learning-based segmentation. All image processing parameters and thresholds were kept consistent across all analyzed samples to ensure comparability of the measurements.

**Supplemental Information 2: Deriving equation (11) for mathematical modeling of pulmonary capillaries**

To estimate the area ratio, $\frac{A_{pleura}}{A_{septa}}$, we approximate the septum geometry as a regular hexagonal lattice, where $L$ is the side length of each hexagon and $\tau$ is the thickness of each line segment. As such, each unit of the lattice consists of a larger hexagon with side length $L$ that circumscribes a smaller hexagon with side length $l<L$, where the smaller hexagon represents the airspace. The total area of the lattice, $A_{pleura}$, is then proportional to the area of the larger hexagon, while the total area of the airspace, $A_{airspace}$, is proportional to the area of the circumscribed hexagon. Consequently, the total area of the septa, $A_{septa}$, is proportional to the difference of these two areas. Thus, from the equation for the area of a regular hexagon, we have equations *A(1a-c)*.

$$\begin{aligned} A_{pleura}=\frac{3\sqrt{3}}{2}L^{2}\#A\left( 1a \right) \end{aligned}$$

$$\begin{aligned} A_{airspace}=\frac{3\sqrt{3}}{2}l^{2}\#A\left( 1b \right) \end{aligned}$$

$$\begin{aligned} A_{septa}=\frac{3\sqrt{3}}{2}{(L}^{2}-l^{2})\#A\left( 1c \right) \end{aligned}$$

The ratio $\frac{A_{septa}}{A_{pleura}}$ immediately follows from equations *A(1a)* and *A(1c)*, yielding *A(2)*.

$$\begin{aligned} \frac{A_{septa}}{A_{pleura}}=1-\left( \frac{l}{L} \right)^{2}\#A\left( 2 \right) \end{aligned}$$

If the wall thickness $\tau$ is given, then by simple trigonometry, the length of the circumscribed hexagon is given by *A(3).*

$$\begin{aligned} l=L-\frac{\sqrt{3}}{3}\tau\#A\left( 3 \right) \end{aligned}$$

Substituting *A(3)* into *A(2)*, we recover *A(4)*.

$$\begin{aligned} \frac{A_{septa}}{A_{pleura}}=1-\left( \frac{L-\frac{\sqrt{3}}{3}\tau}{L} \right)^{2}=\frac{2\sqrt{3}}{3}\left( \frac{\tau}{L} \right)-\left( \frac{\tau}{L} \right)^{2}\#A\left( 4 \right) \end{aligned}$$

When the second term is much smaller than the first, as in the case of the lung’s geometry, equation *A(4)* approximately equals equation *A(5)*.

$$\begin{aligned} \frac{A_{septa}}{A_{pleura}}\approx\frac{2\sqrt{3}}{3}\left( \frac{\tau}{L} \right)\#A\left( 5 \right) \end{aligned}$$

Inverting *A(5)* and substituting $\tau=2R$, we recover *A(6)*, which reproduces equation (11).

$$\begin{aligned} \frac{A_{pleura}}{A_{septa}}\approx\frac{\sqrt{3}}{4}\left( \frac{L}{R} \right)\#A\left( 5 \right) \end{aligned}$$

**References:**

1. Kirillov, A., Mintun, E., Ravi, N., Mao, H., Rolland, C., Gustafson, L., … Girshick, R. (2023). Segment Anything. arXiv:2304. 02643.
